# Chromosome-level hybrid *de novo* genome assemblies as an attainable option for non-model organisms

**DOI:** 10.1101/748228

**Authors:** Coline C. Jaworski, Carson W. Allan, Luciano M. Matzkin

## Abstract

The emergence of third generation sequencing (3GS; long-reads) is making closer the goal of chromosome-size fragments in *de novo* genome assemblies. This allows the exploration of new and broader questions on genome evolution for a number of non-model organisms. However, long-read technologies result in higher sequencing error rates and therefore impose an elevated cost of sufficient coverage to achieve high enough quality. In this context, hybrid assemblies, combining short-reads and long-reads provide an alternative efficient and cost-effective approach to generate *de novo*, chromosome-level genome assemblies. The array of available software programs for hybrid genome assembly, sequence correction and manipulation is constantly being expanded and improved. This makes it difficult for non-experts to find efficient, fast and tractable computational solutions for genome assembly, especially in the case of non-model organisms lacking a reference genome or one from a closely related species. In this study, we review and test the most recent pipelines for hybrid assemblies, comparing the model organism *Drosophila melanogaster* to a non-model cactophilic *Drosophila*, *D. mojavensis*. We show that it is possible to achieve excellent contiguity on this non-model organism using the DBG2OLC pipeline.

## 1. Introduction

Whole genome sequencing is a major target in evolutionary biology, because it provides the material to study how a species’ genome evolves. Notably, whole genome data provides the opportunity to study recombination and large rearrangement events, differential molecular evolution across the genome, and imprints of selection throughout the genome, ultimately improving our knowledge of how species evolve and diverge (Ellegren 2014, Rudman et al., 2018). To increase our understanding of such evolutionary processes, we need to expand the range of studied organisms to non-model organisms, for which the access to well resolved genome assemblies is often lacking.

Thanks to third generation sequencing (3GS) from platforms such as PacBio (Rhoads & Au 2015) and Nanopore (Urban, Bliss, Lawrence & Gerbi 2015), *de novo* genome assemblies of non-model organisms can be obtained, but one drawback from such technologies is the high error rate. *De novo* hybrid assemblies combine long-reads and short-reads (Illumina technology; Bentley et al., 2008) to achieve high contiguity and accuracy while reducing sequencing costs through lower coverage of long-reads data (Ye, Hill, Wu, Ruan & Ma, 2016).

There is a constantly increasing panel of tools to assemble reads and polish genome assemblies. Identifying the pipeline most optimized to one’s needs is one obstacle, and applying it to the actual data is another one, especially in the absence of bioinformatic expertise, since guidelines and practical implementations remain limited. In addition, many of those pipelines are not tested on non-model organisms and assume that the samples are from model organisms where extreme inbreeding and high homozygosity is commonly feasible. In the present study, we reviewed the most recent whole genome assembly pipelines, and selected a promising pipeline relying on hybrid technology (Chakraborty, Baldwin-Brown, Long & Emerson, 2016). We tested it thoroughly with the aim of an optimized assembly, using DNA data from both *D. melanogaster* as a model species, and *D. mojavensis* from the Sonora, Mexico population as a non-model species. Ultimately, this new *D. mojavensis* assembly from Sonora will be used in a much larger upcoming genomic study using *de novo* assemblies of multiple cactophilic species and populations (Matzkin, unpublished data). We provide here an analysis of the effects of different parameters on the quality of the final assembly, assessed by a combination of universal tools (contigs length and N50 as a measure of contiguity; BUSCO score as a measure of quality and completeness (Waterhouse et al., 2017) and a reference-based tool, Quast (Gurevich, Saveliev, Vyahhi & Tesler, 2013) which compares the assembly to a reference genome. We show a significant improvement of assembly quality compared with results from Chakraborty et al. (2016) simply by tuning parameters and we provide guide parameters for assemblies with similar coverage of non-model organism DNA. Finally, we tested the pipeline on *D. mojavensis* from the Santa Catalina Island, California population using Nanopore long-read data instead of PacBio data.

## 2. Materials and Methods

### 2.1 *Drosophila mojavensis* sequencing

We used flies from the *Drosophila mojavensis* isofemale line MJ 122 collected from Guaymas, Sonora Mexico in 1998, thereafter SON. Prior to DNA extraction the flies were raised on banana-molasses medium (Coleman, Benowitz, Jost & Matzkin, 2018) with 125 µg/ml ampicillin and 12.5 µg/ml tetracycline to reduce bacteria contamination of the sequencing data. The sequencing methods for short-read data (paired ends and mate pairs) have been described in Allan & Matzkin (2019). Sequencing technologies and coverage for the different data sets are summarized in Table 1.

**Table 1.** Sequencing technology and coverage of each data set.

| Organism | Genome size | Sequencing technology | Data output | Coverage |
| --- | --- | --- | --- | --- |
| *Drosophila melanogaster* | ~ 140 Mb | <u>Illumina</u> HiSeq 2000 | 16.8 Gb<sup>a</sup> | 120x |
|  |  | <u>PacBio</u> RS II | 16.1 Gb<sup>b</sup> | 115x |
| *Drosophila mojavensis* (Sonora) | ~ 160 Mb | <u>Illumina</u> HiSeq 2500<br>100 bp paired ends | 10.1 Gb<sup>a</sup> | 63x |
|  |  | 2,500 bp mate pairs | 5.9 Gb<sup>c</sup> | 37x |
|  |  | <u>PacBio</u> Sequel | 10.4 Gb<sup>b</sup> | 65x |
| *Drosophila mojavensis* (Catalina) | ~ 160 Mb | <u>Illumina</u> NextSeq 500<br>150 bp paired ends | 9.1 Gb<sup>a</sup> | 55x |
|  |  | Oxford <u>Nanopore</u> MinION | 15.2 Gb | 95x |

<sup>a</sup> calculated as the total number of bases in data files after trimming from *Platanus\_trim*.
<sup>b</sup> calculated as the total number of bases in data files obtained from converting .bax.h5 files into .bam files, and converting the obtained subreads.bam files into fasta files, with no specific trimming
<sup>c</sup> calculated as the total number of bases in data files after trimming from *Platanus\_internal\_trim*.

#### 2.1.2 DNA extraction for PacBio Sequencing – Protocol optimization

Due to the long-read potential of PacBio Sequencing Systems, extra care must be taken during extraction to produce high molecule weight DNA. Attempts at using both QIAGEN DNeasy Blood &Tissue Kit and QIAGEN MagAttract HMW DNA Kit failed to produce sufficiently long strands of DNA. As such, a chloroform-based extraction method was used. This relatively simple method is low cost and the only specialized equipment needed is a refrigerated centrifuge. To consistently recover enough DNA for two PacBio libraries, 150 flies of each sex were used for extraction. Flies were starved for two hours in groups of 50 per vial and then frozen at −80° C in 1.5 mL tubes. A lysis solution containing Tris HCI buffer 0.1M pH 8.0, EDTA 0.1 M pH 8.0, and 1% SDS was prepared and stored at room temperature to prevent the SDS from precipitating. While on ice, 500 µL of lysis solution was added to each tube of flies, followed by 2.5 µL of QIAGEN Proteinase K to reduce DNA degradation. Using a plastic pestle, flies in each tube were hand-homogenized by gently grinding them. Hand homogenization resulted in slightly lower amounts of DNA recovered, however the size of DNA fragments were longer compared to when using a battery-operated pestle motor to homogenize. The mixture was incubated at 65 °C for 30 min with gentle mixing halfway through. To further reduce DNA fragmentation, tubes were cooled to 37 °C for three minutes and another 2.5 µL of QIAGEN Proteinase K was added. Tubes were incubated for an additional 30 min at 37 °C. After incubation, 70 µL of 4 M potassium acetate was added, mixed by inversion, and then placed on ice to incubate for 30 minutes. In a 4 °C Eppendorf 5920R centrifuge, the tubes were spun for 30 min at 18,000 rcf to pull debris to the bottom of the tubes. For each tube, the supernatant was transferred to new tubes avoiding as much debris as possible. One volume of chloroform:isoamyl alcohol 24:1 was added to each tube and gently inverted 40 times, and then centrifuged at 4 °C for 5 min at 10,500 rcf. The upper phase was transferred to a new tube while being careful to not disturb the interface. The DNA was precipitated by adding 350 µL of 2-propanol and gently inverting the tube. At this point visible threads of DNA were apparent. To pellet the precipitate, the tubes were centrifuged at 4 °C for 5 min at 10,500 rcf. The supernatant was discarded and the pellet was washed with 1 mL of room-temperature 70% ethanol. The tube was inverted to insure washing of the pellet and tube. A final 4 °C centrifugation for 2 min at 10,500 rcf was performed. The ethanol was removed with a pipettor as completely as possible and the pellet dried for 10-15 min in a fume hood. 30 µL of Tris-EDTA pH 8.0 was added to each tube to resuspend the DNA. While the pellet normally goes into solution relatively easily, it can be placed at 4 °C overnight to insure resuspension. The six tubes were then combined to a single tube and 3 µL of QIAGEN RNaseA was added and incubated for 30 min at 37 °C. The DNA was delivered as this resuspended solution for PacBio sequencing.

#### 2.1.3 PacBio Sequencing

PacBio sequencing was performed at the Arizona Genomics Institute (Tucson, AZ, U.S.A.). DNA was sized in a 1% agarose pulsed field gel electrophoresed at 1-50 sec linear ramp, 6 volts/cm, 14 °C in 0.5X TBE buffer for 20 hours (BioRad). The marker used was a lambda ladder Midrange PFG I (New England Biolabs). The resultant DNA smear had a large mass in the 35-65 kb range (Fig. 1). DNA purity was verified using a NanoDrop One Microvolume UV Spectrophotometer with ratios 260/280 and 260/230 over 1.8. Quantity (150 ng/µL in 180 µL = 27 µg) was determined by a Qubit Fluorometer (Life Technologies), and was consistently lower than that measured with the Nanodrop. PacBio sequencing libraries were prepared from 6 µg starting material each, following manufacturers protocol for a 20 Kb Template Preparation Using BluePippin™ Size-Selection System (www.pacb.com). The library was size-selected, on a BluePippin, at 20 kb using high pass with S1 Marker (Sage Sciences). The final library was damage-repaired, bead-purified and quantified. Sequencing was performed on a PacBio Sequel instrument following manufacturer’s instructions. The sequencing primer annealed was v3, the sequencing kit was v2.1. Two libraries were loaded on two separate SMRT cells with magbeads at concentrations of 25 pmol and 35 pmol, respectively. Sequencing was carried out for collection of 10 hr movies on 1 M SMRT cells.

**Fig. 1.**
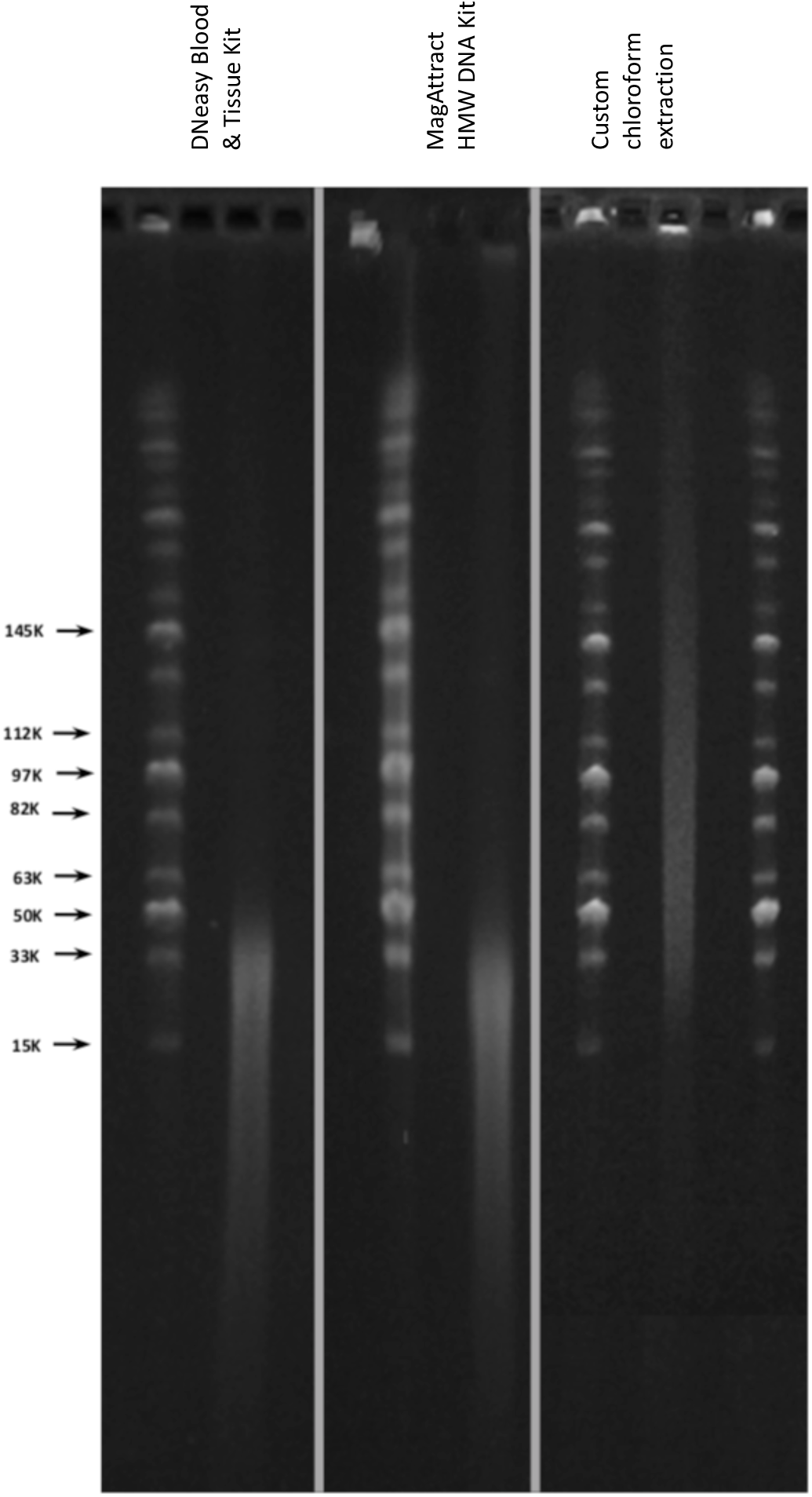
Size profile of DNA from *D. mojavensis* (Sonora) extracted with three different methods. Images of three gels, corresponding to each method, have been collated here, using the same ladder (sizes shown on the left).

### 2.2. *Drosophila melanogaster* and *D. mojavensis* from Catalina public sequencing data

To generate the *D. melanogaster* assembly (hereafter, Dmel), PacBio data was retrieved from the NCBI Short-read Archive SRX499318 (Kim et al., 2014). This data set contained 42 PacBio RS II SMRT cells from male *D. melanogaster* ISO1 flies. We used data from 20 randomly selected cells only to obtain a coverage similar to our data sets (cell numbers SRR1204085, SRR1204088, SRR1204451, SRR1204466, SRR1204467, SRR1204469, SRR1204471, SRR1204472, SRR1204473, SRR1204481, SRR1204482, SRR1204485, SRR1204486, SRR1204615, SRR1204617, SRR1204690, SRR1204691, SRR1204692, SRR1204693, and SRR1204696). We used the SMRT Illumina HiSeq 2000 100 bp paired-end data from male *D. melanogaster* ISO1 flies, which was retrieved from the European Nucleotide Archive ERX645969 (Miller, Smith, Hawley & Bergmann, 2013).

For the *D. mojavensis* assembly from the Santa Catalina Island, California population (hereafter, CAT), Nanopore sequencing data was kindly provided by Miller, Staber, Zeitlinger & Hawley (2018). Short-read Illumina data was retrieved from the NCBI Short-read Archive SRR6425997 (Miller et al., 2018).

### 2.3. Computing resources

All the programs were run on the UA Research Computing High Performance Computing (HPC) at the University of Arizona. The cluster used is composed of 28 core processors with 168 gb RAM per node, and is run via a PBS-Pro grid system. All the programs used were installed under a user python virtual environment (pip). The majority of the programs used are available as Bioconda packages for easy installation in non-cluster environments (Grünings et al., 2018). They are also provided as Docker containers through Bioconda which can be run through *Singularity* (https://sylabs.io/) on cluster systems. All command lines are provided in Appendix.

### 2.4. Assembly pipelines

#### 2.4.1. DBG2OLC Pipeline

The DBG2OLC Pipeline is composed of three main steps: (i) the hybrid assembly via the DBG2OLC program, (ii) the long-read assembly only, and (iii) the merging of those two assemblies (Fig. 2).

**Fig. 2.**
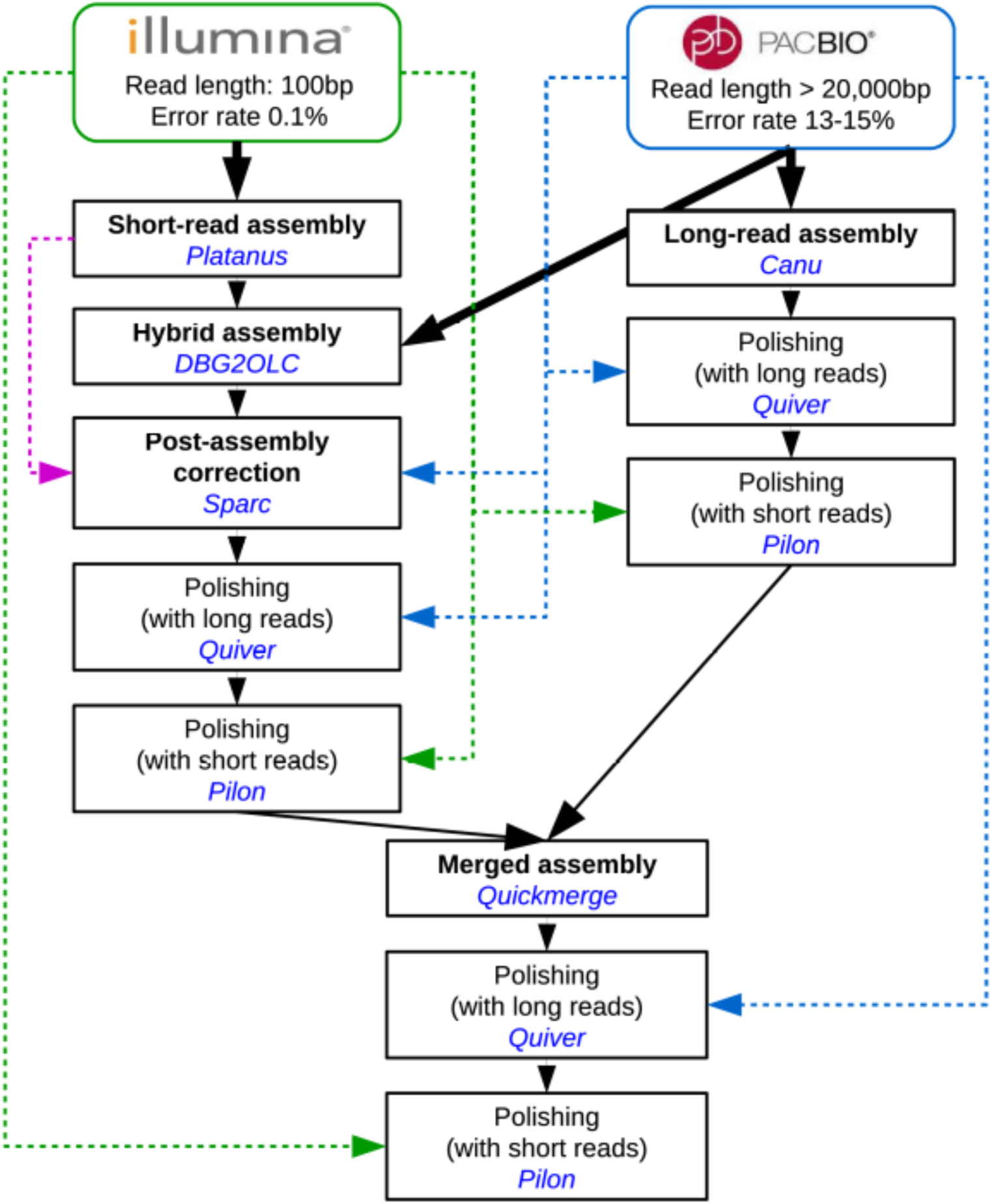
DBG2OLC Pipeline, including the final merging step and the polishing steps.

##### (i) Hybrid assembly

*DBG2OLC* uses contigs from a short-read assembly and maps them to the raw long-reads, which are then compressed into the list of the short-read’s contig identifiers (Ye et al., 2016). A best overlap graph is constructed from those compressed long-reads before uncompressing them into a consensus sequence. This method is both highly accurate and extremely fast (Ye et al., 2016). Then, the consensus contigs, or backbones are corrected using *Sparc* (Ye & Ma, 2016). *Sparc* builds a sparse k-mer graph (k-mers in different positions are treated independently) using the contigs identifiers’ list associated with each raw long-read. All short-read contigs are then aligned to their associated long-read using the *Blasr* aligner from the PacBio SMRT toolkit (SMRT Link v4.0.0), previously *Pbdagcon*, which is the most time-consuming step. *Sparc* finally uses these alignments to refine the graph and create a polished consensus sequence. In the present study, we tested two competing short-read assemblers, *SparseAssembler* (provided with the DBG2OLC installation package) (Ye, Ma, Cannon, Pop & Yu, 2012), and *Platanus* version 1.2.4 (Kajitani et al., 2014). We used the March 2019 version of *DBG2OLC* (Ye et al., 2016), the January 2015 version of *Sparc* (Ye & Ma, 2016), and *Blasr* 5.3.5 (b30da0) (SMRT Link v4.0.0). Note that we began working with an older version of *Blasr* which was signficantly slower and led to slightly different results. For this reason, and because programs often include third party packages, it is important to keep track of each version used and physically separate the repositories, so the SMRT toolkit was installed in an independent directory with no direct link to the user bin, except for the *Blasr* program. We modified the split_and_run_sparc.sh script available from the *Sparc* Github repository so as to call the split_reads_by_backbone.py script externally (Appendix I), and to set the number of chores used by *Blasr* from the command line. This way, it is easier to rerun the time-consuming Sparc step in case of crash from where it stopped, and after moving the already corrected backbones into another directory.

The hybrid assembly was then polished using the PacBio tool in the SMRT toolkit (SMRT Link v4.0.0). The version of the PacBio correction tool is frequently updated along with chemistry technology of PacBio sequencing, therefore the version *Quiver* (v2.1.0) was used for *D. melanogaster* (sequenced in 2014 on a PacBio RS II system; Kim et al., 2014), and the version *Arrow* (v2.1.0) was used for *D. mojavensis* (sequenced in 2017 on a PacBio Sequel system installed with SMRT Link v4.0.0, see above). For simplicity, we will thereafter refer to that step simply as ‘*Quiver*’. *Quiver* aligns the raw PacBio reads to the assembled and corrected contigs output by *Sparc*, and uses a consensus caller to polish them (Chin et al., 2013). Lastly, the hybrid assembly was polished using *Pilon* v1.22 (Walker et al., 2014). *Pilon* uses raw short-reads aligned to the assembly with the *Bowtie2* aligner version 2.2.9 (Langmead, Trapnell, Pop & Salzberg, 2009), to first find and correct SNPs and small indels (base error consensus), and secondly local misassemblies (alignment discrepancies scan) that are reassembled using paired ends and mate pairs (if provided). Parameters were optimized at each step: (a) choice of the short-read assembler (*Platanus* vs. *SparseAssembler* with kmer-size 39 or 53); (b) *DBG2OLC* parameters, based on recommended optimization ranges (Ye et al., 2016): MinOverlap in [20; 150]; AdaptiveTh in [0.002; 0.02]; KmerCovTh in [2; 10] and MinLen in [200; 2,000]; default values were otherwise used (k = 17; LD1 = 0); (c) ContigTh 0 (default) vs. 1 (recommended for > 100x PacBio coverage only); and (d) *Sparc* one vs. two iterations. These parameters are summarized in Table 2.

**Table 2.** *DBG2OLC* parameters.

| Assembly | Assembler | MinOverlap | AdaptiveTh | MinLen | KmerCovTh | Number of<br>*Sparc*<br>iterations | ContigTh |
| --- | --- | --- | --- | --- | --- | --- | --- |
| P0 | *Platanus* | 20 | 0.002 | 200 | 2 | 2 | 1 |
| P1 | *Platanus* | 20 | 0.020 | 200 | 2 | 2 | 1 |
| P2 | *Platanus* | 100 | 0.020 | 200 | 2 | 2 | 1 |
| P3 | *Platanus* | 150 | 0.002 | 200 | 2 | 2 | 1 |
| S3 | *SparseAssembler* | 150 | 0.002 | 200 | 2 | 2 | 1 |
| P4 | *Platanus* | 150 | 0.010 | 200 | 2 | 2 | 1 |
| P4f | *Platanus* | 150 | 0.010 | 200 | 2 | 3 | 1 |
| P4g | *Platanus* | 150 | 0.010 | 200 | 2 | 3 | 2 |
| P5 | *Platanus* | 150 | 0.015 | 200 | 2 | 2 | 1 |
| P6 | *Platanus* | 150 | 0.020 | 200 | 2 | 2 | 1 |
| P6f | *Platanus* | 150 | 0.020 | 200 | 2 | 3 | 1 |
| P6g | *Platanus* | 150 | 0.020 | 200 | 2 | 3 | 2 |
| S6 | *SparseAssembler* | 150 | 0.010 | 200 | 2 | 2 | 1 |
| P4a | *Platanus* | 150 | 0.010 | 2000 | 2 | 2 | 1 |
| P5a | *Platanus* | 150 | 0.015 | 2000 | 2 | 2 | 1 |
| P6a | *Platanus* | 150 | 0.020 | 2000 | 2 | 2 | 1 |
| P6b | *Platanus* | 150 | 0.020 | 5000 | 2 | 2 | 1 |
| P4y | *Platanus* | 150 | 0.010 | 200 | 10 | 2 | 1 |
| P6y | *Platanus* | 150 | 0.020 | 200 | 10 | 2 | 1 |
| P6x | *Platanus* | 150 | 0.020 | 200 | 5 | 2 | 1 |

##### (ii) Long-read assembly only

The long-read assembly was created using *Canu* v1.5 (Koren et al., 2017), which significantly outperforms its older version *Celera Assembler* (PbcR) used in Chakraborty et al. (2016) as well as other assemblers, notably by using an adaptive kmer weighting which both improves the efficiency and the quality of the assembly of highly repetitive genomic regions. We tested two parameters for the correctedErrorRate: 0.039 vs. 0.055 (low end and middle value of recommended range, Table 3, Koren et al., 2017). Note that this adjustment is limited by coverage, and thus intrinsic to the analyzed data set. We ran the three *Canu* steps (correction, trimming and assembly) separately using the options -correct, -trim and -assemble (see Appendix) to optimize the assembly step without running again the first two steps. Similarly to the hybrid assembly, the long-read only assembly was polished using both *Quiver* and *Pilon*.

**Table 3.** *Canu* and *Quickmerge* parameters.

| Species | Assembly | *Canu*<br>correctedErrorRate | Merge parameters |  |  |
| --- | --- | --- | --- | --- | --- |
|  |  |  | Query assembly | l (bp) | lm (bp) |
| Dmel | P4 | 0.039 | Hybrid | N50: 10,000,000 | 10,000 |
|  | P4f | 0.039 | Hybrid | N50: 10,000,000 | 10,000 |
|  | P4n | 0.039 | Hybrid | N50/2: 5,000,000 | 10,000 |
|  | P4o | 0.039 | Hybrid | N50/4: 2,500,000 | 10,000 |
|  | P4o'' | 0.039 | Hybrid | 1,000,000 | 10,000 |
|  | P4fo'' | 0.039 | Hybrid | 1,000,000 | 10,000 |
|  | P4p | 0.039 | Hybrid | N50: 10,000,000 | 0 |
|  | P4q | 0.039 | Hybrid | 0 | 0 |
|  | P4r | 0.055 | Hybrid | 0 | 0 |
|  | P4s | 0.039 | Long-read | 0 | 0 |
| SON | P6 | 0.039 | Hybrid | N50: 2,600,000 | 10,000 |
|  | P6f | 0.039 | Hybrid | N50: 2,600,000 | 10,000 |
|  | P6n | 0.039 | Hybrid | N50/2: 1,300,000 | 10,000 |
|  | P6n'' | 0.039 | Hybrid | 1,000,000 | 10,000 |
|  | P6fn'' | 0.039 | Hybrid | 1,000,000 | 10,000 |
|  | P6o | 0.039 | Hybrid | N50/4: 0,650,000 | 10,000 |
|  | P6p | 0.039 | Hybrid | N50: 2,600,000 | 0 |
|  | P6q | 0.039 | Hybrid | 0 | 0 |
|  | P6r | 0.055 | Hybrid | 0 | 0 |
|  | P6s | 0.039 | Long-read | 0 | 0 |
| CAT | P6r'' | 0.055 | Hybrid | 1,000,000 | 10,000 |
|  | P6fr'' | 0.055 | Hybrid | 1,000,000 | 10,000 |

##### (iii) Assembly merging

The hybrid assembly and long-read only assembly were merged after polishing using the *Quickmerge* tool v0.2 (Chakraborty et al. 2016). *Quickmerge* uses *MUMmer* (v 3.0) (Kurtz et al. 2004) to align the two assemblies and find the unique best alignment (using the -delta-filter option in *MUMmer*). *Quickmerge* then identifies high confidence overlaps between the two assemblies to find seed contigs (i.e., contigs that can be extended at both ends). Finally, it merges the overlapping contigs using sequences from the reference (donor assembly) into the query (acceptor assembly). The optimization consisted of trying both the hybrid and long-read assemblies as reference vs. query, and varying the l and lm cutoff parameters in *Quickmerge* (Table 3). Lastly, the merged assembly was polished using *Quiver* and *Pilon*.

#### 2.4.2. Test of the *DBG2OLC* pipeline with Nanopore long-reads

We ran the DBG2OLC pipeline on CAT sequencing data using Nanopore reads instead of PacBio reads for the long-read only assembly, the hybrid assembly, and the polishing steps. We used the optimal parameter set (2.4.1; P6), except for *Canu*, for which we had to increase the correctedErrorRate to 0.055 to recover 97% of the genome, while we could recover only 51% using a correctedErrorRate of 0.039. Instead of *Quiver*, we used *Nanopolish* version 0.11.0 (Simpson et al., 2017). Similar to *Quiver* and *Pilon*, raw Nanopore reads were first aligned to the target assembly using the *Bwa* aligner version 0.7.17 (Li & Durbin, 2010). *Nanopolish* then generates an improved consensus sequence.

#### 2.4.3. Alternative Pipelines

*DBG2OLC* was identified as the only pipeline, among the most recently published assemblers, allowing the assembly of long-reads prior to correction. Alternatively, long-reads may be corrected prior to assembly, as is the case in the *Canu* pipeline. Other possible correction tools include *LSCplu*s (Hu, Sun & Sun, 2016), a modified version of the *MHAP* tool (indexing kmers used to build the assembly graph in *Celera*; (Carvalho, Dupim & Goldstein, 2016), *HALC* (Bao & Lan, 2017), and *FMLRC* (Holt, Wang, Jones & McMillan, 2016), and most recently (not tested here) *MECAT* (Xiao et al., 2017) and *Jabba* (Miclotte et al., 2016). The *LSCplus* package was not available at the time of our study, and we were therefore not able to assess its efficiency. The modified *MHAP* tool was implemented in *Celera* only (the older version of *Canu*), which thus yielded poor results in term of assembly contiguity, and this solution was abandoned. Note however that some aspects of kmers indexing as proposed in the modified *MHAP* tool have now been implemented in *Canu* (Koren et al., 2017), and were therefore implicitly used in our *DBG2OLC* Pipeline. *FMLRC*, although it correctly performed the long-read correction, proved to be non-compatible with *Canu* (Holt et al., 2016). Therefore, this alternative was abandoned as well.

*HALC* corrects long-reads by (i) aligning them to the contigs from a short-read assembly, (ii) constructing a graph from this alignment, and (iii) finding the best path in the graph to correct each long-read. It relies on *Blasr* (SMRT Link v4.0.0) for the alignment and on *LoRDEC* (Salmela & Rivals 2014) for the correction. We used the one version of *HALC* available, *Blasr* 5.3.5 and *LoRDEC* 0.6 with the *GATB* library 1.0.6. After read correction with *HALC*, we ran *Canu* (- assemble option) with a correctedErrorRate = 0.039. The contiguity of the assembly was orders of magnitude worse than when using the *Canu* correction tool and the same assembly parameters (*HALC* correction: N50 = 488,850; total length = 138,021,997; max length = 2874227; *Canu* pipeline: N50 = 10,9990,654; total length = 151,043,692; max length = 25,950,142). This might have been improved by parameter optimization of both the *HALC* correction step and the assembly step with the *Canu* assembler, but due to the strong difference in contiguity we chose to not utilize *HALC*. Therefore, we focused on optimizing the *DBG2OLC* pipeline only.

### 2.5. Assembly quality check

Comparisons between assemblies and quality assessment were performed based on assembly statistics from *Quast* version 4.6.2 (Gurevich et al., 2013) by comparing each assembly to a reference genome to estimate the number of global and local misassemblies as well as the number of mismatches and indels. For both general statistics (number of fragments, N50) and error rates (presented in Tables 4-6), we used contigs longer than 400 bp only, so as to run the program faster. We also calculated *BUSCO* scores using the diptera (odb9) set of Benchmarking Universal Single-Copy Orthologs (Waterhouse et al. 2017). We used the reference genomes FB2017_01and FB2015_02 (Consortium DG, 2007) released on FlyBase (Thurmond et al., 2019) for Dmel and CAT, respectively. For SON, we used a template assembly constructed based on the Catalina reference genome (Allan & Matzkin, 2019). For each data set, we extracted only the fragments that have been previously designated to chromosomes (i.e., for Dmel, the four chromosomes; and for SON and CAT the 39 biggest scaffolds), so as to run quality assessment faster. We are aware that using a template assembly as a reference for SON may introduce biases especially in terms of number of misassemblies, due to the evolutionary history of the *D. mojavensis* populations (Matzkin, 2014) therefore the results must be considered carefully. However, this provides a valid guide to make relative comparisons between assemblies created here. *Quast* relies on *MUMmer* v3.23 (nucmer aligner v3.1; Kurtz et al., 2004) to align the assembly to the reference genome, and includes metrics and methods from the *GAGE* assessment tool (Salzberg et al., 2012) and other tools.

**Table 4.**
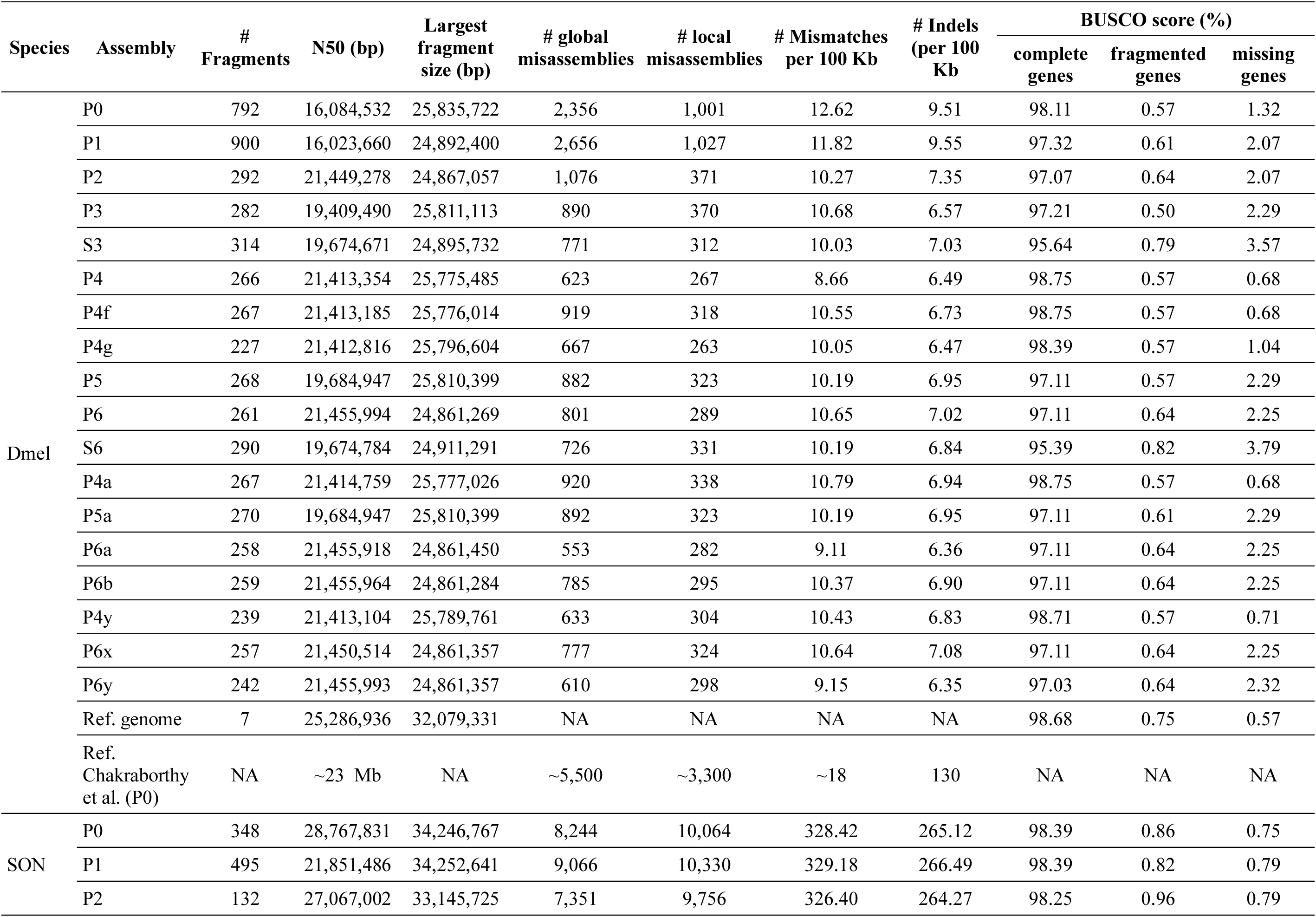
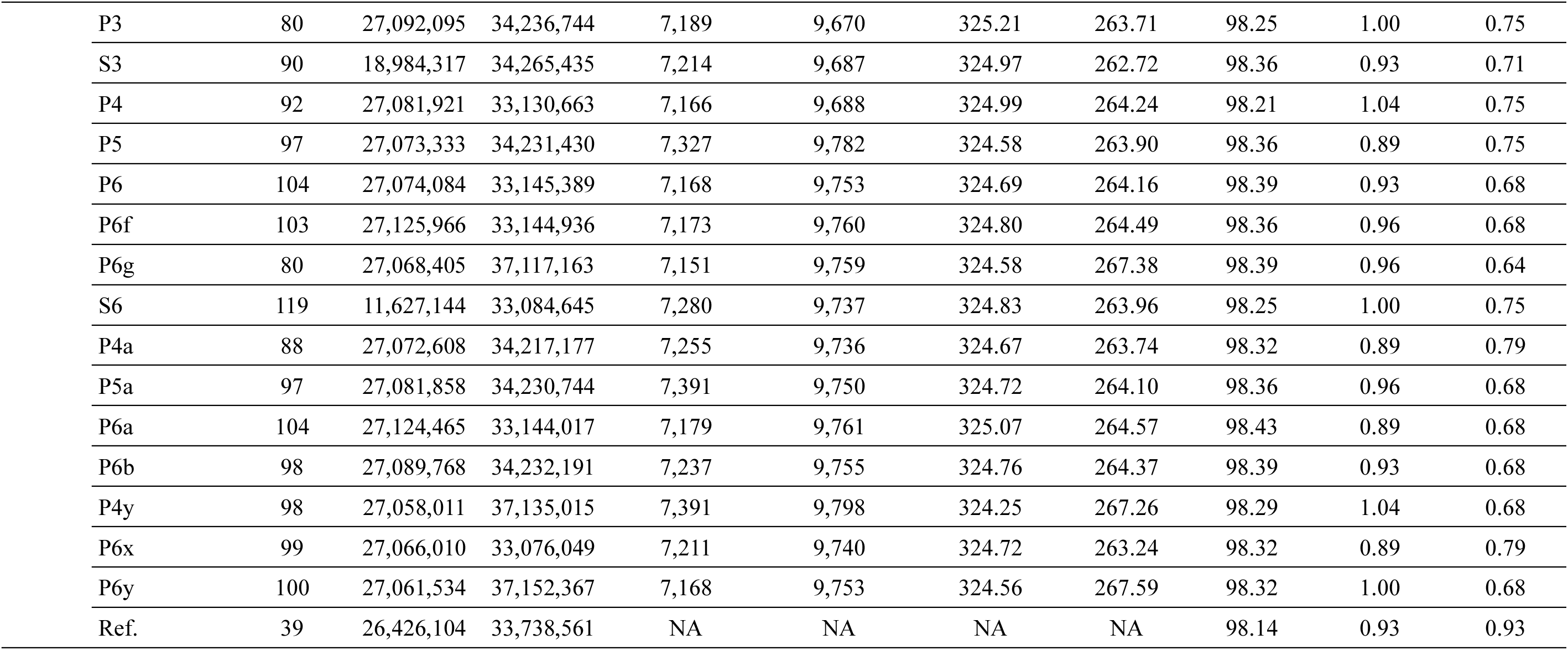
*DBG2OLC* parameter optimization: contiguity and accuracy. Assemblies refer to parameter sets defined in Table 2.

**Table 5.** *Canu* and *Quickmerge* parameter optimization: contiguity and accuracy. Assemblies refer to parameter sets defined in Table 3.

| Species | Assembly | # Fragments | N50 (bp) | Largest fragment size (bp) | # global misassemblies | # local misassemblies | # Mismatches per 100 Kb | # Indels per 100 Kb | BUSCO score |  |  |
| --- | --- | --- | --- | --- | --- | --- | --- | --- | --- | --- | --- |
|  |  |  |  |  |  |  |  |  | complete genes | fragmented genes | missing genes |
| Dmel | P4 | 266 | 21,413,354 | 25,775,485 | 623 | 267 | 8.66 | 6.49 | 98.75 | 0.57 | 0.68 |
|  | P4f | 267 | 21,413,185 | 25,776,014 | 919 | 318 | 10.55 | 6.73 | 98.75 | 0.57 | 0.68 |
|  | P4o | 265 | 21,413,352 | 25,775,437 | 635 | 275 | 8.57 | 6.42 | 98.75 | 0.57 | 0.68 |
|  | P4o'' | 210 | 21,413,344 | 25,775,300 | 638 | 295 | 8.10 | 6.02 | 98.79 | 0.57 | 0.64 |
|  | P4fo'' | 211 | 21,413,185 | 25,776,408 | 943 | 340 | 9.90 | 6.20 | 98.79 | 0.57 | 0.64 |
|  | P4q | 116 | 21,413,360 | 25,775,323 | 1,561 | 521 | 8.72 | 5.77 | 98.79 | 0.57 | 0.64 |
|  | P4r | 113 | 21,450,483 | 25,801,485 | 1,751 | 582 | 8.66 | 5.75 | 98.61 | 0.75 | 0.64 |
|  | P4s | 338 | 14,528,003 | 25,770,616 | 3,684 | 1,145 | 12.34 | 7.34 | 98.79 | 0.57 | 0.64 |
|  | Ref. | 7 | 25,286,936 | 32,079,331 | NA | NA | NA | NA | 98.68 | 0.75 | 0.57 |
| SON | P6 | 104 | 27,074,084 | 33,145,389 | 7,168 | 9,753 | 324.69 | 264.16 | 98.39 | 0.93 | 0.68 |
|  | P6f | 103 | 27,125,966 | 33,144,936 | 7,173 | 9,760 | 324.80 | 264.49 | 98.36 | 0.96 | 0.68 |
|  | P6n | 88 | 27,074,290 | 33,145,104 | 7,357 | 9,788 | 324.44 | 263.58 | 98.39 | 0.93 | 0.68 |
|  | P6n'' | 67 | 27,074,601 | 33,145,127 | 7,261 | 9,728 | 325.29 | 263.80 | 98.29 | 0.96 | 0.75 |
|  | P6fn'' | 66 | 27,125,795 | 33,144,975 | 7,271 | 9,727 | 325.23 | 263.97 | 98.25 | 1.00 | 0.75 |
|  | P6o | 65 | 27,074,498 | 33,145,149 | 7,261 | 9,718 | 325.17 | 263.76 | 98.32 | 0.93 | 0.75 |
|  | P6q | 62 | 27,074,467 | 33,145,189 | 7,470 | 9,752 | 325.83 | 263.63 | 98.32 | 0.93 | 0.75 |
|  | P6r | 58 | 27,027,841 | 34,117,449 | 7,245 | 9,165 | 329.97 | 262.71 | 93.25 | 0.93 | 5.82 |
|  | P6s | 151 | 27,122,727 | 34,181,614 | 9,159 | 10,205 | 328.59 | 263.06 | 98.14 | 0.93 | 0.93 |
|  | Ref. | 39 | 26,426,104 | 33,738,561 | NA | NA | NA | NA | 98.14 | 0.93 | 0.9 |
| CAT | P6n'' | 79 | 12,454,906 | 23,097,599 | 2,272 | 3,104 | 56.41 | 67.39 | 98.39 | 1.00 | 0.61 |
|  | P6fn'' | 79 | 12,457,238 | 23,102,169 | 2,274 | 2,949 | 56.40 | 65.70 | 98.64 | 0.82 | 0.54 |
|  | Ref. | 39 | 26,866,924 | 34,148,556 | NA | NA | NA | NA | 98.11 | 0.93 | 0.96 |
|  | Ref. Miller et al. (2018) | 122 | 5.0 Mb | NA | NA | NA | 0.22 | 0.052 | 98 | NA | NA |

**Table 6.** Improvement of contiguity and quality throughout the pipeline. Abbreviations: Sr: short-read assembly; H: hybrid assembly; Lr: long-read assembly; M: merged assembly; Q: Quiver polishing; P: Pilon polishing. Here MQP corresponds to P4 for Dmel and P6 for SON (Tables 4, 5).

| Species | Assembly | # Fragments | N50 (bp) | Largest fragment (bp) | # global misassemblies | # local misassemblies | # Mismatches per 100 kb | # Indels per 100 kb | BUSCO score |  |  |
| --- | --- | --- | --- | --- | --- | --- | --- | --- | --- | --- | --- |
|  |  |  |  |  |  |  |  |  | complete genes | fragmented genes | missing genes |
| Dmel | Sr | 15,404 | 22,245 | 250,600 | 138 | 22 | 3.05 | 0.68 | 96.96 | 2.07 | 0.96 |
|  | H | 302 | 5,369,803 | 20,351,387 | 449 | 391 | 28.34 | 570.44 | 66.02 | 15.51 | 18.47 |
|  | HQ | 302 | 5,378,161 | 20,387,636 | 685 | 439 | 8.26 | 13.51 | 95.03 | 2.00 | 2.97 |
|  | HQP | 302 | 5,378,529 | 20,385,575 | 689 | 464 | 8.91 | 7.78 | 97.89 | 0.57 | 1.54 |
|  | Lr | 426 | 10,086,116 | 24,845,957 | 3,935 | 2,126 | 16.25 | 12.16 | 89.03 | 6.50 | 4.47 |
|  | LrQP | 426 | 10,090,934 | 24,862,005 | 4,002 | 2,131 | 12.23 | 8.07 | 98.21 | 0.57 | 1.21 |
|  | M | 266 | 21,413,390 | 25,776,101 | 603 | 288 | 8.95 | 7.19 | 98.75 | 0.57 | 0.68 |
|  | MQP | 266 | 21,413,354 | 25,775,485 | 623 | 267 | 8.66 | 6.49 | 98.75 | 0.57 | 0.68 |
|  | Ref. | 7 | 25,286,936 | 32,079,331 | NA | NA | NA | NA | 98.68 | 0.75 | 0.57 |
| SON | Sr | 57,046 | 3,385 | 45,376 | 346 | 2,539 | 219.24 | 200.08 | 89.10 | 7.93 | 2.97 |
|  | H | 136 | 9,893,295 | 18,894,064 | 6,773 | 9,558 | 331.40 | 457.22 | 88.14 | 8.40 | 3.47 |
|  | HQ | 136 | 9,840,048 | 18,808,757 | 6,898 | 9,646 | 324.72 | 271.29 | 98.54 | 0.93 | 0.54 |
|  | HQP | 136 | 9,834,752 | 18,808,038 | 6,893 | 9,663 | 324.81 | 265.21 | 98.68 | 0.82 | 0.50 |
|  | Lr | 343 | 2,678,315 | 8,942,850 | 8,790 | 9,912 | 332.66 | 316.57 | 96.57 | 1.93 | 1.50 |
|  | LrQP | 343 | 2,679,816 | 8,945,441 | 9,055 | 10,035 | 328.02 | 262.04 | 98.14 | 1.04 | 0.82 |
|  | M | 104 | 27,077,180 | 33,146,112 | 7,167 | 9,792 | 325.14 | 264.44 | 98.43 | 0.89 | 0.68 |
|  | MQP | 104 | 27,074,084 | 33,145,389 | 7,168 | 9,753 | 324.69 | 264.16 | 98.39 | 0.93 | 0.68 |
|  | Ref. | 39 | 26,426,104 | 33,738,561 | NA | NA | NA | NA | 98.14 | 0.93 | 0.9 |

Finally, assemblies were aligned to their reference genome using MUMmer4 (Marçais et al., 2018) and plotted against the reference genome using Dot (https://github.com/dnanexus/dot).

### 2.6. Test of the *DBG2OLC* pipeline with Nanopore long-reads

We tested the DBG2OLC pipeline on *D. mojavensis* population Catalina using Nanopore long-reads instead of PacBio long-reads with parameters optimized for SON: *Platanus* short-read assembler; *DBG2OLC* parameters MinOverlap 150; AdaptiveTh 0.020; KmerCov 2; MinLen 200; number of *Sparc* iterations 2 or 3 (both tested: P6r” vs. P6fr”. For the *Canu* assembly, we used the correctedErrorRate = 0.055 since lower rates resulted in incomplete genome (0.039: 73.7 %; 0.045: 86.7 %; 0.055: 93.8 %). For the merged assembly, we used parameters as optimized for SON: the hybrid assembly as query; l = 1 Mb and lm = 10,000 bp (Table 3).

## 3. Results and Discussion

### 3.1. DNA preparation

The custom chloroform extraction led to a remarkable increase in the sizes of DNA fragments (light band between 30 and 120 Kb, right panel, Fig. 1) compared with standard extraction kits (left and middle panels, Fig. 1) for which the majority of fragments were shorter than 30 Kb. Long fragments in DNA libraries significantly increase DNA quantity output by PacBio sequencing (www.pacb.com).

### 3.2. Optimization of the DBG2OLC Pipeline

#### 3.2.1. Short-read assembler

*Platanus* and *SparseAssembler* with a kmer size of 53 bp resulted in very similar assemblies; *SparseAssembler* with a kmer size of 39 bp led to reduced contiguity; and applying two successive rounds of *SparseAssembler* 53 bp-Kmer size did not improve the short-read assembly. In the final merged assemblies, the use of *SparseAssembler* always led to slight decrease in contiguity (comparing P3 to S3 and P6 to S6 for both Dmel and SON, Table 4). *SparseAssembler* slightly reduced error rates but also BUSCO scores for Dmel, with limited effects for SON (Fig. 3B). We also observed that differences in P6 and S6 for SON mainly resided in highly repetitive regions. Since *Platanus* is especially recommended for genomes with high level of heterozygosity, which are more likely in non-model organisms, we used *Platanus* for all the subsequent assemblies in this study.

**Fig. 3.**
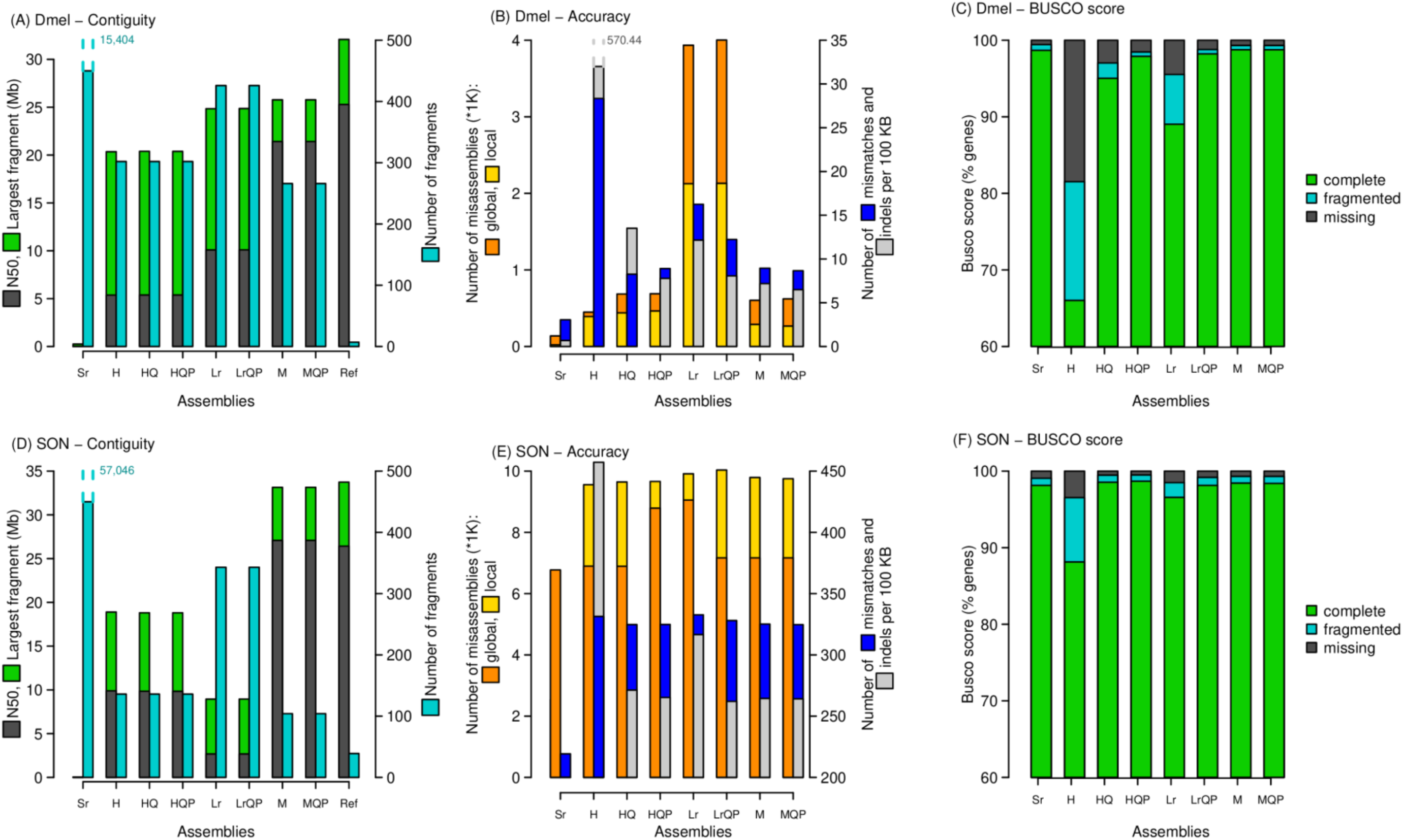
Contiguity (A, D), error level (B, E) and Busco score (C, F) for Dmel (A-C) and SON (D-F) assemblies, at each step of the pipeline. Significantly larger values are printed above dashed lines. Assembly parameters are described in Table 6.

#### 3.2.2. DBG2OLC parameters

We varied the DBG2OLC parameters MinOverlap, AdaptiveTh, KmerCovTh and MinLen to simultaneously optimize the contiguity and quality of the final assembly. Misassemblies created during the first steps of the hybrid assemblies were overall not resolved later, which makes that step key to the optimization. P0 corresponds to the reference set of parameters used in Chakraborty et al. (2016). Note that P0 has similar contiguity as shown in Chakraborty et al. (2016) for the assembly with ∼100x coverage although with lower error rates, indicating that our analysis using their parameter set resulted in a very similar result.

MinOverlap had a major effect on final assemblies, with a major improvement of contiguity (reduced number of fragments, increased N50, increased length of longest fragment; Fig. 3A,C) and of accuracy (reduced number of global and local misassemblies, reduced number of mismatches and indels; Fig. 3B, 3D) as seen in the P0 vs. P3 and P1 vs. P6 comparisons. This came at a cost of a slight decrease in BUSCO score for Dmel but not SON. Only an increase of MinOverlap up to 150 (the maximum recommended value for more than 50x coverage of PacBio reads) led to an optimal lower number of misassemblies (P2 vs. P6).

AdaptiveTh had little influence, except when MinOverlap was kept low: it decreased contiguity and accuracy (P2 vs. P0). For assemblies with high MinOverlap, we found that P3 was less fragmented than P4, P5 or P6 for SON and P4 was the least fragmented for Dmel. P6 was the best compromise between contiguity and accuracy for SON, with the highest BUSCO score. P4 was the best compromise and with the highest BUSCO score for Dmel. Although coverage in both Illumina short-reads and PacBio long-reads was lower in SON than Dmel (Table 1), the quality of PacBio long-reads was higher (longer reads thanks to DNA extraction protocol, and more recent PacBio technology), which might have facilitated the better results with the more stringent AdaptiveTh. Therefore, for PacBio data sequenced with the newest technology and coverage higher than 50x, we recommend to use the most stringent value for AdaptiveTh, of 0.020.

High KmerCovTh values resulted in major global misassemblies in SON (assessed with Mummer plots, not shown), with the largest fragment longer than the theoretical longest fragment in the Reference assembly (P4y vs. P4 and P6y,x vs. P6). It also caused a slight increase in error rates and a slight decrease in BUSCO score. In Dmel, no major global misassembly was detected, however error rates were higher and BUSCO scores slightly lower. We recommend to use KmerCovTh = 2, especially when using high AdaptiveTh. Using ContigTh = 1 had similar effects (major global misassemblies; P6g vs. P6 for SON) than high KmerCovTh values, so we recommend not using it.

Increased MinLen from 200 to 2,000 resulted in a slight increase in contiguity for both Dmel and SON (PX vs. PXa). Error rates and BUSCO scores were not notably different, unless MinLen was increased up to 5,000 in which too many reads were parsed out, leading to higher error rates. Due to the absence of major effects and according to DBG2OLC recommendations (Ye et al. 2016), we recommend to not increase MinLen above 200, because there is a risk to parse out under-represented genome regions.

Increasing the number of *Sparc* iterations from 2 to 3 allowed a higher contiguity of large fragments, although with little effect on overall statistics. Increasing *Sparc* iterations is supposed to reduce the number of chimeras. Since it had no negative effect, we recommend using three iterations.

Based on our tests, we recommend the following DBG2OLC parameters, for 50x or more of PacBio reads from recent technology: MinOverlap 150; AdaptiveTh 0.020; KmerCov 2; MinLen 200; number of *Sparc* iterations 3. These parameters were used throughout the next optimization step (except AdaptiveTh 0.010 for Dmel).

#### 3.2.3. Canu parameters

Increasing correctedErrorRate from 0.039 to 0.055 slightly increased contiguity of the merged assembly (P4q vs. P4r for Dmel and P6q vs. P6r for SON; Table 5). However, it also increased error rates overall, and decreased the BUSCO complete genes score, especially for SON. We therefore recommend to keep the *Canu* correctedErrorRate as low as possible, to 0.039.

#### 3.2.4. Quickmerge parameters

Parameters used in *Quickmerge* are shown in Table 3, and results in Table 5. Using the long-read assembly as the Query assembly resulted in a strong decrease in contiguity compared with the opposite (P4s vs. P4q for Dmel and P6s vs. P6q for SON). It also considerably increased error rates for both species, and slightly decreased BUSCO complete gene score for SON. Similarly to Chakraborty et al. (2016), we therefore recommend to use the hybrid assembly as query.

We also tested the impact of the l and lm parameters. Using low lm with high l resulted in identical assemblies (P4p vs. P4 for Dmel and P6p vs. P6 for SON) since backbones were already parsed out due to high lm. Also, using lm = N50 or l = N50/2 resulted in identical assemblies for Dmel. Otherwise, decreasing l resulted in lower number of fragments but higher error rates. However, using a too high lm value would prevent smaller fragments from being merged, this is why we recommend an intermediate value of 1 Mb (P6fn”).

#### 3.2.5. Polishing

Polishing with both Quiver/Arrow and Pilon did not affect the contiguity (number of fragments, N50, and largest fragment; Table 6, Fig. 3) for either species. Conversely, it significantly reduced the number of indels on hybrid, long-read and merged assemblies. The number of mismatches was also reduced to a lesser extent. One drawback was the increase in number of misassemblies, except for the merged assembly. Finally, polishing increased the BUSCO score, especially on the hybrid assembly.

### 3.3. Test of the *DBG2OLC* pipeline with Nanopore long-reads

Compared with Miller et al. (2018), the CAT merged assembly was more contiguous and with a higher BUSCO score, but with higher error rates (Table 5, P6r” vs. Ref. Miller), likely due to the multiple polishing steps performed by Miller et al. (2018). Compared with SON, CAT assemblies were less contiguous but with higher BUSCO scores (CAT-P6r” vs. SON-P6n” and CAT-P6fr” vs. SON-P6fn”). These results show that the DBG2OLC pipeline outperforms other ones in terms of contiguity using either PacBio or Nanopore long-read data.

### 3.4. Benefits of using the *DBG2OLC* pipeline and demonstration of effectiveness on a non-model species

By merging hybrid and long-read only assemblies, we considerably increased the contiguity compared with that of the hybrid assembly or the long-read only assembly (Table 6, Fig. 3), as shown in Chakraborty et al. (2016). Also, error rates were lower than the long-read assembly, especially for Dmel. To obtain such low error rates with long-read data only, a higher coverage would have been necessary representing a significant increase in sequencing cost (discussed in Chakraborty et al., 2016). For this study the *D. mojavensis* Illumina sequencing was performed in 2011, if using current sequencing core prices it will total ∼ $178 (PE 150 HiSeq lane ∼$1,300 [only 1/12th of a lane needed for the 160 Mb *D. mojavensis* genome]; quality control of library $15; library preparation $50-$400 [depending if done in-house or by a core]). The PacBio sequencing was performed using a Sequel system, totaling $3,190 (library preparation $495 × 2 libraries; SMRT cells $1,100 × 2). Given the recent release of PacBio’s Sequel II system the cost for a similar amount of long-reads would be approximately ∼$740 (library preparation ∼$450, SMRT cell $1,750 [would only need 1/6^th^ of a cell for *D. mojavensis*]), therefore the *de novo* assembly described in this study could be built for less than $1,000.

One major improvement of the merged assembly (P6fn”) in SON is that the 2q^5^ inversion in the Müller element D (described in Ruiz, Heed & Wasserman, 1990) is now resolved, with the two breakpoints clearly bridging the three chromosome parts (Fig. 4). This was not the case in the hybrid assembly or the long-read only assembly (not shown). The Müller elements B, D and E in our merged assembly P6fn” are in one piece and correspond to 99.24%, 99.11% and 96.67% respectively of the corresponding chromosomes in the CAT reference genome. The Müller element C was composed of three pieces in P6fn” accounting for 99.94% of the length of the Müller element C in the CAT reference, and the Müller element A was more fragmented, as is also the case in the CAT reference genome and all fragment lengths summed up to 94.77% of the total size in the AT reference. However, in the CAT reference genome, D was in two fragments that were joint in our assembly.

**Fig. 4.**
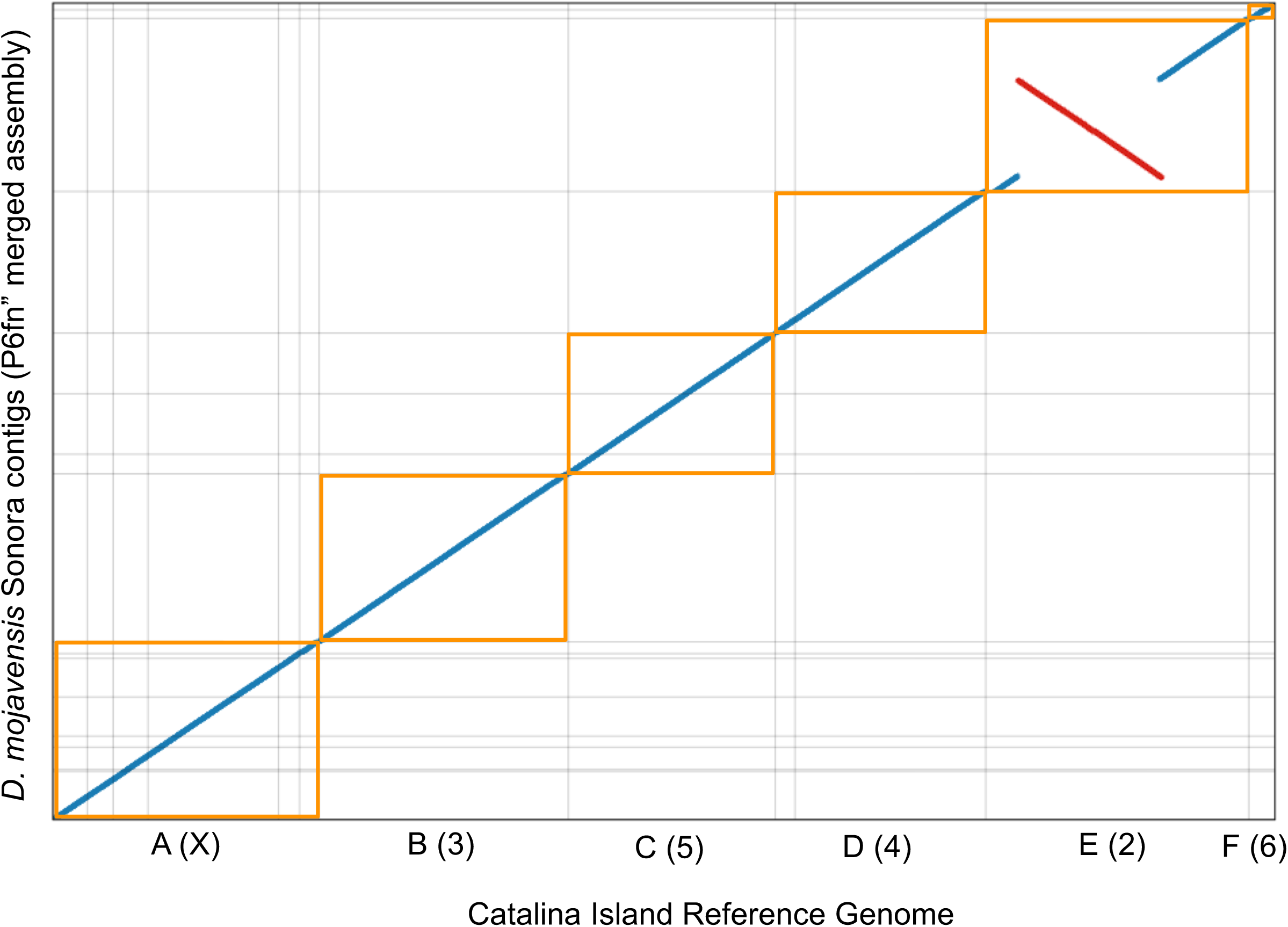
Alignment of SON merged assembly P6fn” (y-axis) on the *D. mojavensis* (Catalina) reference genome (x-axis). Only fragments longer than 900 Kb are shown. Müller elements (chromosomes) of the reference genome (Catalina) are shown. Yellow boxes represent a single Müller element. Gray horizontal lines indicate the contigs from the SON assembly.

## 4. Conclusion

In the not too distant past genomic analysis was limited to just a set of a few model laboratory species. Although this has led to unprecedented advances in our understanding of genetics and genomics, in many instances such studies lacked an ecological context. Genome assemblies of non-model species tended to be more fragmented or tended to be built using a genome from a related model species, which is problematic if interested in trait mapping or genome structure evolution. Current sequencing and computational advancements have liberated our dependence on classical laboratory model species. Here we have outlined a widely applicable computational pipeline and sets of parameters to facilitate the construction of chromosome or nearly-chromosome level genomic assemblies in a non-model species. Overall, our PacBio merged assembly performed better than using Nanopore reads, but more work is still needed to assess any differences across multiple species, especially with newer advances to the sequencing platforms.

## 5. Acknowledgments

We would like to thank Rod Wing, David Kudrna and Jayson Talag at the Arizona Genomics Institute for their assistance in the PacBio sequencing. This work was supported with funding from the National Science Foundation (IOS-1557697) to LMM and the University of Arizona to LMM.

## 7. Data accessibility

Raw reads are available at NCBI’s SRA (https://www.ncbi.nlm.nih.gov/sra) under accession number XXXXXX.

## 8. Author contribution

CCJ, CWA and LMM reviewed the literature and selected the pipeline. CWA performed DNA extraction. CCJ performed the bioinformatics work with the help of CWA. CCJ wrote the manuscript with contributions from LMM and CWA.

## Appendix. Command lines to call each program used in the study.

## (1) Data manipulation

Merge .bam files from several PacBio sequencing cells (node=1:ncpus=1:mem=6gb):

∼/Programs/SMRTtools/smrtcmds/bin/bamtools merge -list ListPBreadsFiles.fofn -out PBreads_all.bam

Merge two .fasta files: (node=1:ncpus=1:mem=6gb):

cat reads1.fasta reads2.fasta > reads1_and_2.fasta

Convert .bax.h5 to .bam (node=1:ncpus=1:mem=6gb):

∼/Programs/SMRTtools/smrtcmds/binbax2bam my_movie.*.bax.h5 -o my_movie

Convert .bam to .fastq (node=1:ncpus=1:mem=6gb):

∼/Programs/SMRTtools/smrtcmds/bin/samtools fastq my_reads.bam > my_reads.fastq

OR

∼/Programs/SMRTtools/smrtcmds/bin/bam2fastq my_reads.bam -o my_reads

Convert .bam to .fasta (node=1:ncpus=1:mem=6gb):

∼/Programs/SMRTtools/smrtcmds/bin/bam2fasta my_reads.bam -o my_reads

Convert .fastq to .fasta with the *prinseq-lite tool* (v0.20.4) (node=1:ncpus=1:mem=6gb):

perl prinseq-lite.pl -fastq my_reads.fastq -out_format 1

Convert .fastq to .fasta with the *FASTX* toolkit (v0.0.13) (node=1:ncpus=1:mem=6gb):

fastq_to_fasta -i my_reads.fastq -o my_reads.fasta

## (2) Short-read assembly

Trimming (with *Platanus* trimmer)

(node=1:ncpus=5:mem=30gb;cput=10:00:00;walltime=02:00:00):

platanus_trim my_PE_reads_1.fastq my_PE_data_2.fastq -t 5

*Platanus* assembler (node=1:ncpus=14:mem=65gb; cput=28:00:00;walltime=02:00:00):

platanus assemble -o Shortread_contigs -f my_PE_reads_[12].fastq.trimmed -t 16 -m 75 2> Plat_SON2.log

*SparseAssembler*, with kmer size 53 (node=1:ncpus=1:mem=6gb; cput=02:00:00; walltime=02:00:00):

SparseAssembler LD 0 k 53 g 15 NodeCovTh 1 EdgeCovTh 0 GS 165000000 p1 my_PE_reads_1.fastq.trimmed p2 my_PE_reads_2.fastq.trimmed

## (3) Hybrid assembly

*DBG2OLC* step (node=1:ncpus=1:mem=6gb; cput=02:50:00:walltime=02:50:00):

DBG2OLC k 17 KmerCovTh 2 MinOverlap 150 AdaptiveTh 0.002 LD1 0 MinLen 200 Contigs Shortread_contigs.fasta RemoveChimera 1 f my_PBreads.fasta

Note: DBG2OLC has memory limitations. If run on node=1:ncpus=2:mem=12gb, it will still use one chore only, but more memory as allowed, and run twice as fast.

Building list of contigs identifiers (node=1:ncpus=1:mem=6gb; cput=00:05:00;walltime=00:05:00) (requires python2):

split_reads_by_backbone.py -b backbone_raw.fasta -o ./Cons_backbones -r Shortread_contigs_and_PBreads.fasta -c DBG2OLC_Consensus_info.txt

*Sparc* step ((node=1:ncpus=28:mem=90gb; cput=84:00:00;walltime=03:00:00):

sh split_and_run_sparc_ncpus_new.sh ./Cons_backbones 2 28 > Sparclog_Part1.txt

## (4) Long-read only assembly with Canu

canu -p PB_Only_Assembly -d /path_to_curr_dirr genomeSize=123m correctedErrorRate=0.039 - useGrid=true -maxThreads=16 -maxMemory=90 -gridEngineThreadsOption=“-l select=1:ncpus=16:mem=100gb” -gridEngineMemoryOption=“-l walltime=02:00:00” -gridOptions=“-W group_list=my_group_ID -q standard” -pacbio-raw /path_to_Pbreads/Pbreads.fasta

Note: Because the *Canu* pipeline calls a master script, more parameters, normally passed to the PBS script have to be sent to the command line when calling *Canu*. The option -nanopore-raw was used when assembling Nanopore reads.

## (5) Assembly merging

Using the *Quickmerge* wrapper, and with hybrid assembly as donor and long-read only assembly as acceptor (node=1:ncpus=1:mem=6gb; cput=00:25:00;walltime=00:25:00; requires python/3):

merge_wrapper.py Hybrid_assembly.fasta longread_assembly.fasta -l 10000000 -lm 10000

## (6) Quiver polishing

Align PB reads (in .bam format) to assembly with *Pbalign* (node=1:ncpus=28:mem=168gb; cput=224:00:00;walltime=08:00:00):

∼/Programs/SMRTtools/smrtcmds/bin/pbalign --nproc 28 my_PBreads.bam my_assembly.fasta aligned_PBreads.bam

Index aligned PB reads (node=1:ncpus=1:mem=6gb; cput=00:10:00;walltime=00:10:00):

∼/Programs/SMRTtools/smrtcmds/bin/pbindex aligned_PBreads.bam

Index assembly (node=1:ncpus=1:mem=6gb; cput=00:02:00;walltime=00:02:00):

∼/Programs/SMRTtools/smrtcmds/bin/samtools faidx my_assembly.fasta

Run Quiver (node=1:ncpus=28:mem=168gb; cput=154:00:00;walltime=05:30:00):

∼/Programs/SMRTtools/smrtcmds/bin/quiver -j 28 -r my_assembly.fasta -o my_assembly_polished.fasta aligned_PBreads.bam

Run *Arrow* (node=1:ncpus=28:mem=168gb; cput=154:00:00;walltime=05:30:00):

∼/Programs/SMRTtools/smrtcmds/bin/arrow -j 28 -r my_assembly.fasta -o my_assembly_polished.fasta aligned_PBreads.bam

## (7) Pilon polishing

Index the assembly file (node=1:ncpus=6:mem=6gb; cput=00:05:00;walltime=00:05:00):

bowtie2-build my_assembly.fasta my_assembly

Align short-reads to assembly with Bowtie2 (node=1:ncpus=28:mem=168gb; cput=42:00:00;walltime=01:30:00):

bowtie2 -x my_assembly -1 my_PE_reads_1.fastq.trimmed -2 my_PE_reads_2.fastq.trimmed -S Aligned_reads.sam -p 28

Convert .sam to .bam (node=1:ncpus=1:mem=6gb; cput=00:10:00;walltime=00:10:00):

samtools view -bS Aligned_reads.sam > Aligned_reads.bam

Sort reads (node=1:ncpus=28:mem=168gb; cput=07:00:00;walltime=00:15:00):

samtools sort Aligned_reads.bam -o Aligned_reads_sorted.bam -@ 28

Index reads (node=1:ncpus=6:mem=6gb; cput=00:02:00;walltime=00:02:00):

samtools index Aligned_reads_sorted.bam -@ 28

Run *Pilon* with paired ends only (node=1:ncpus=28:mem=168gb; cput=14:00:00;walltime=00:30:00; requires java/8):

java -Xmx64G -jar pilon-1.22.jar --genome my_assembly.fasta --frags Aligned_PEreads_sorted.bam -- output ./my_assembly_polished --threads 28

Run *Pilon* with paired ends and mate pairs

(node=1:ncpus=28:mem=168gbcput=28:00:00;walltime=01:00:00; requires java/8):

java -Xmx64G -jar pilon-1.22.jar --genome my_assembly.fasta --frags Aligned_PEreads_sorted.bam -- jumps Aligned_MPreads_sorted.bam--output ./my_assembly_polished --threads 14

Note: *Pilon* seems to try to use more cpus than allowed with the --thread option and requires more memory to run on both paired ends and mate pairs. To counter this problem, we actually ran it on 28 cpus but passed less cpus (just enough to avoid memory issue) to the --thread option.

## (8) Nanopolish polishing

Index the raw Nanopore reads (node=1:ncpus=1:mem=6gb; cput=02:30:00;walltime=02:30:00):

nanopolish index -d ./path_to_Fast5 -f List_summary.fofn -v ./Reads_basecalled_pass.fastq

Index the assembly file (node=1:ncpus=6:mem=6gb; cput=00:10:00;walltime=00:10:00):

bwa index my_assembly.fasta

Align short-reads to assembly with *Bwa* (node=1:ncpus=28:mem=168gb; cput=161:00:00;walltime=05:45:00):

bwa mem -x ont2d -t 28 my_assembly.fasta /path_to_raw_reads/Reads_basecalled_pass.fastq | samtools sort -o Reads.sorted.bam -@ 28

samtools index Reads.sorted.bam -@ 28

Run *Nanopolish* (node=1:ncpus=28:mem=168gb; cput=854:00:00;walltime=30:30:00):

python nanopolish_makerange.py my_assembly.fasta | cat > Chunks.txt

more Chunks.txt | parallel --results ./nanopol.res -P 7 nanopolish variants --consensus --faster -o polished.{1}.vcf -w {1} -r ./Reads_basecalled_pass.fastq -b ./Reads.sorted.bam -g ./my_assembly.fasta -t 4 --min-candidate-frequency 0.2

Note: *Nanopolish* was unable to handle paths to directory others than the working directory, therefore we placed both raw indexed reads, the draft assembly and the sorted reads in the same directory.

Convert *.vcf to .fasta* (node=1:ncpus=1:mem=6gb; cput=00:03:00;walltime=00:30:00):

nanopolish vcf2fasta -g ./my_assembly.fasta ./polished.*.vcf > ./my_assembly_polished.fasta

## (9) Quality assessment

*Quast* (node=1:ncpus=5:mem=30gb; cput=05:00:00;walltime=01:00:00;requires python/3)

∼/Programs/quast-4.6.2/quast.py -o /./ -R /path_to_reference_genome/Reference.fasta --threads 5 --min-alignment 400 --no-plot --no-html --no-icarus --labels My_assembly --eukaryote /path_to_assembly/My_assembly.fasta

Note: walltime will depend on how divergent is the draft assembly from the reference assembly

*Busco* (node=1:ncpus=14:mem=84gb; cput=25:40:00;walltime=01:50:00; requires python/3, augustus/3 and hmmer/3)

python ∼/Programs/busco/scripts/run_BUSCO.py -i /path_to_assembly/My_assembly.fasta -o output_directory -l ∼/Programs/busco/diptera_odb9/ -m genome -c 14 -sp fly --blast_single_core

## (10) Visualization with Dot

(node=1:ncpus=1:mem=6gb; cput=00:08:00;walltime=00:08:00)

nucmer -c 100 -t 3 -prefix=my_assembly /path_to_reference/Reference_genome.fasta /path_to_assembly/My_assembly_100KB.fasta

python DotPrep.py --delta my_assembly.delta --out outputname --unique-length 10000 --overview 500000

